# Streamlined production, purification, and comparison of recombinant extracellular polyhydroxybutyrate depolymerases

**DOI:** 10.1101/700252

**Authors:** Diana I. Martínez-Tobón, Brennan Waters, Anastasia L. Elias, Dominic Sauvageau

## Abstract

Heterologous production of extracellular polyhydroxybutyrate (PHB) depolymerases (PhaZs) has been of interest for over 30 years, but implementation is sometimes difficult and can limit the scope of research. With the constant development of tools to improve recombinant protein production in *Escherichia coli*, we propose a method that takes characteristics of PhaZs from different bacterial strains into account. Recombinant His-tagged versions of PhaZs (rPhaZ) from *Comamonas testosteroni* 31A, *Cupriavidus* sp., *Marinobacter algicola* DG893, *Pseudomonas stutzeri*, and *Ralstonia* sp. were successfully produced with varying expression, solubility, and purity levels. PhaZs from *C. testosteroni* and *P. stutzeri* were more amenable to heterologous expression in all aspects; however, strategies were developed to circumvent low expression and purity for the other PhaZs. Degradation activity of the rPhaZs was compared using a simple PHB plate-based method, adapted to test for various pH and temperatures. rPhaZ from *M. algicola* presented the highest activity at 15 °C, and rPhaZs from *Cupriavidus* sp. and *Ralstonia* sp. had the highest activity at pH 5.4. The methods proposed herein can be used to test the production of soluble recombinant PhaZs, and to perform preliminary evaluation for applications that require PHB degradation.

## 1 Introduction

The study of extracellular polyhydroxybutyrate (PHB) depolymerases (PhaZs) produced by a variety of microorganisms [1-3] remains an important and evolving research area. Their enzymatic activity results in the degradation of PHB, a natural biodegradable polymer with the potential to replace some currently widely used petroleum-based plastics [4] that increasingly accumulate in the environment [5].

Recombinant protein production is a powerful tool that allows the production of higher levels of proteins in expression systems such as *E. coli*. Optimized recombinant technologies facilitate purification, the study of proteins in isolation, the conception of a platform to modify and improve them, and the development of new applications. In the case of PhaZs, such applications include biosensors — such as time-temperature indicators [6] and pathogen detection platforms [7] — and recycling of biodegradable polymers [8].

Examples of expression of rPhaZs in *E. coli* include PhaZ2–PhaZ3 [9] and PhaZ7 (although for this specific PhaZ better expression was achieved with in *Bacillus subtilis* WB800) [10] from *Pseudomonas lemoignei*, and PhaZ from *Caldimonas manganoxidans* [8,11]. In some cases, purification of rPhaZs has also been performed: several PhaZs from *P. lemoignei* (PhaZ1–PhaZ5) [12,13], PhaZ7 and related mutants [14]), *Pseudomonas stuzeri* [15], *Alcaligenes faecalis* AE122 [16], *Marinobacter* sp. NK-1 [17], *Bacillus megaterium* N-18-25-9 [18], *Pseudomonas mendocina* DSWY0601 [19], and from *Cupriavidus sp.* (formerly *Alcaligenes faecalis* T1) and related mutants [20-24]. However, these studies each required the development of specific methods for heterologous expression of specific PhaZs. In addition, in many cases affinity tags were not employed, requiring significant additional steps for purification [12,13,16,21-23]. These factors impede on the rapidity and scope of studies, even limiting comparisons between PhaZs.

In this study, we established a platform for the rapid expression and purification of extracellular rPhaZs. This was demonstrated with five extracellular PhaZs displaying different properties and of various bacterial origins. Predicted solubility and disulfide bonds (necessary for maintaining proper conformation and activity in many proteins [25]) of the rPhaZs produced were important criteria in selecting the *E. coli* system, specifically the plasmid vector and expression strains. A single platform with simple strategies was successfully employed for expression, purification, and preliminary comparison of degradation performance under different conditions.

## 2 Materials and methods

### 2.1 Bacterial strains and growth conditions

The bacterial strains used for isolation of the PhaZs, cloning and expression, as well as their growth medium and conditions can be found in Table 1. Cell growth was monitored by measuring optical density of the cultures at 600 nm (OD_600_) using a UV-Vis spectrophotometer (Biochrom, Ultrospec 50). Plating was performed on 1.5% w/v agar supplemented with the medium of interest and plates were incubated in a temperature-controlled incubator (Isotemp 500 Series, Fisher Scientific).

**Table 1.**
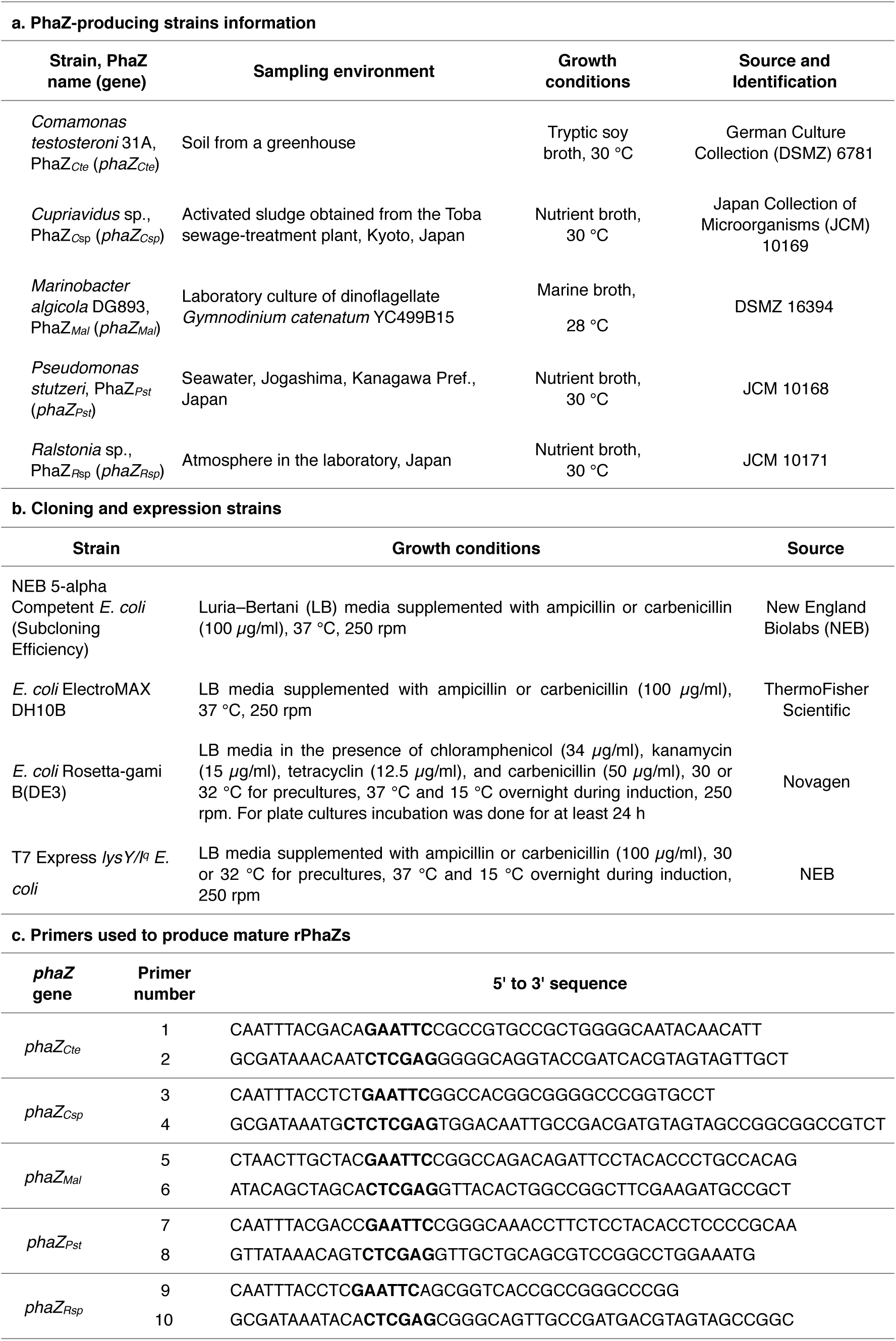
Bacterial strains, conditions, and primers. Growth conditions and information of (a) PhaZ-producing strains and (b) cloning and expression strains; (c) rPhaZs primers.

### 2.2 PhaZs constructs

Genomic DNA was extracted from the PhaZ-producing strains (RNA/DNA purification kit, Norgen Biotek for *C. testosteroni*, and GeneJET Genomic DNA Purification Kit, ThermoFisher Scientific for other strains). Inserts were obtained by amplifications of the mature *phaZ* genes (without signal peptides) — with primers (5’ to 3’ direction, Table 1c) designed according to the GenBank sequences [26] adding restriction sites *Eco*RI and *Xho*I (boldface in Table 1c) through polymerase chain reaction (PCR) (T100 Thermal Cycler, Bio-Rad) using Phusion High-Fidelity DNA Polymerase (ThermoFisher Scientific). All PCR products were purified with QIAquick PCR Purification Kit (Qiagen).

Restriction digestions — using *Eco*RI and *Xho*I (NEB) — and ligations — using T4 DNA ligase (ThermoFisher Scientific) — were performed to obtain PhaZs inserts and to incorporate them into the pET-22b(+) vector (Novagen). This plasmid includes an N-terminal peIB leader sequence for periplasm localization of the protein, and an optional C-terminal His-tag (included in the constructs). Transformation was done chemically with heat shock at 42 °C for 30 s or by electroporation with 0.1 cm gap cuvettes at 1.8 kV for 1 s (Gene Pulser, Bio-Rad), depending on the competent cells required. This was followed by addition of 250 *µ*l of SOC medium (Super Optimal broth with Catabolite repression), incubation for 1 h at 37 °C and 250 rpm, and plating. The constructs were extracted with QIAprep Spin Miniprep Kit (Qiagen) and verified by DNA gel electrophoresis and sequencing (ABI 3730 DNA sequencer, Applied Biosystems). Constructs were inserted in the expression strain *E. coli* Rosetta-gami B(DE3), and, for PhaZ_*Cte*_ and PhaZ_*Mal*_, also in T7 Express *lysY/I*^*q*^ *E. coli*.

### 2.3 Induction screening and His-tag verification

Starter cultures from single colonies of *E. coli* Rosetta-gami B(DE3) or T7 Express *lysY/I*^*q*^ *E. coli* containing the constructs were grown in 5 ml LB with corresponding antibiotics (Table 1b) until reaching an OD_600_ ∼ 0.5 (overnight incubation at ≈ 30 °C and 250 rpm is recommended). 15-ml cultures in LB with antibiotics were inoculated with 1 ml of starter cultures and incubated at 37 °C until reaching an OD_600_ ≈ 0.6. Cultures were separated into 3-ml aliquots, induced with isopropyl-β-d-thiogalactopyranoside (IPTG) (concentrations ranging between 0.01 and 1 mM), and incubated overnight at 15 °C (for PhaZ_*Cte*_ and PhaZ_*Mal*_ incubations at 37 °C for 2 or 4 h were also tested). After incubation, 2 ml of induced cultures were centrifuged (10,000 × *g* and 4 °C for 10 min) and the pellets were placed at -20 °C. B-PER II Bacterial Protein Extraction Reagent (2X) (ThermoFisher Scientific), supplemented with lysozyme (1 mg/ ml, Sigma-Aldrich) and DNAse I (5 units/ml, ThermoFisher Scientific) was used to obtain soluble fractions (SF) (150 *µ*l/pellet), followed by centrifugation at 21,130 × *g* and 4 °C for 30 min. The same procedure was performed with cells carrying empty pET-22b(+) vector and with samples before induction.

SF and insoluble fractions (IF) were then characterized by sodium dodecyl sulfate polyacrylamide gel electrophoresis (SDS-PAGE) analysis, based on the methods described by Laemmli[27]. Briefly, 2X Laemmli sample buffer (Bio-Rad) was added to samples and boiled at 100 °C for 10 min. Loading volumes in the gel were normalized according to OD_600_ of the samples. A Broad-Range protein standard ladder (6.5-210 kD) (Bio-Rad) was used as reference. Samples were run in 12% polyacrylamide gels (Bio-Rad) for 40 min at constant voltage (200 V). The gels were washed three times with Milli-Q water for 10 min and stained with PageBlue protein staining solution (ThermoFisher Scientific) for 1 h under gentle agitation. The gels were then washed with Milli-Q water. Images of the gels were acquired under UV exposure (AlphaImager EC, Alpha Innotech) or with a regular camera. In the case of PhaZ_*Cte*_ and Phaz_*Mal*_, the presence of the His-tag was verified through Western blot analysis by using mouse anti-His6 monoclonal antibody, and goat anti-mouse DyLight 488 secondary antibody (Life Sciences).

### 2.4 Expression and purification of PhaZs

30 mL of transformed *E. coli* Rosetta-gami B(DE3) at OD_600_ of 0.5 were added to 1 L LB with antibiotics. Cultures were grown at 37 °C for approximately 5 h, until OD_600_ reached ≈ 0.6. IPTG was added (0.05 and 1 mM for PhaZ_*Mal*_, and 1 mM for all other PhaZs), and the cultures were incubated overnight at 15 °C for expression. Cultures were then centrifuged at 10,000 × *g* and 4 °C for 10 min, and the pellets were placed at -20 °C. Protein extraction was performed on thawed pellets using 5 ml of B-PER II mixture with Halt™ Protease Inhibitor Cocktail, EDTA-Free (100X) (ThermoFisher Scientific) to obtain SF containing PhaZs.

Purification was performed at 4 °C using His GraviTrap columns (GE Healthcare). Equilibration was done with 10 ml of B-PER II before extracted soluble fractions were applied to the column, followed by a wash with 10 ml of binding buffer (50 mM sodium phosphate, 500 mM NaCl, pH 7.4). All three solutions contained 20 mM imidazole. His-tagged rPhaZs were eluted with 3 ml of elution buffer (20 mM sodium phosphate, 500 mM NaCl, pH 7.4, with 150 mM imidazole for PhaZ_*Mal*_ and 500 mM for all other PhaZs). 1-mL aliquots with 50% glycerol were stored at -20 °C. Purified rPhaZs were verified through SDS-PAGE and quantified with Bradford Protein Assay (microassay procedure, Bio-Rad) using bovine serum albumin as standard. Amicon Ultra 0.5-mL filters (Millipore) were used for PhaZ_*C*sp_, PhaZ_*Mal*_, and PhaZ_*R*sp_, which required further purification.

Buffer exchange was done prior to assays in which imidazole and glycerol caused interference with Amicon Ultra 0.5-mL filters or dialysis (Slide-A-Lyzer MINI Dialysis Devices, 20K MWCO, Thermo Scientific).

### 2.5 PHB plates rPhaZs activity comparison

Rapid PHB degradation assays were performed by dispensing 100 *µ*l of soluble fractions in cylindrical wells made in double-layer mineral medium/agar plates containing PHB (medium 474: 20 ml first layer — mineral medium with agar (0.016 g/ ml) — and 10 ml second layer — mineral medium with agar supplemented with 0.66 ml of sterile PHB suspension). The plates were pierced to produce cylindrical wells for the deposition of samples. Plates were then incubated at 30 °C in a temperature-controlled incubator (Isotemp 500 Series, Fisher Scientific). PHB degradation was assessed by the presence of halos.

The effects of pH and temperature on degradation of PHB by rPhaZs were investigated. Since the PHB plates had pH 7.00, experiments at pH 4.27 and 5.35 were performed using mineral medium with sodium acetate and acetic acid buffer solutions at the desired pH (both solutions at 0.2 M) [28]. The agar and low pH buffer were autoclaved separately. Each rPhaZ was diluted to a concentration of ≈ 2 *µ*g/ml, and 100 *µ*l were deposited for each condition in duplicates. Plates were incubated at 15 °C and 37 °C for pH 7.00, and at 37 °C for pH 4.27 and 5.35. Pictures were taken over four weeks of incubation. Degradation was assessed by the formation of transparent halos on the PHB plates and measuring the halo diameters (minus well diameter) using ImageJ 1.46r (National Institutes of Health, USA).

## 3 Results and Discussion

### 3.1 Selection of expression platform

An *E. coli*-based recombinant protein production system was selected, based on its relative success to produce rPhaZs [9,11-13,15-19] and the wide commercial offer of vectors and hosts. The sequences of mature PhaZs (based on respective references) were processed using a solubility predictor (PROSO II, [29,30]), and the theoretical isoelectric points (pI) and molecular weights (Mw) were calculated using the Compute pI/Mw tool from the ExPASy Bioformatics Resources Portal (SIB) [31-33] (Table 2). Since PhaZ_*Cte*_ was classified as insoluble (predicted solubility score 0.503), and the other PhaZs had scores (0.657– 0.765) near the PROSO II threshold for solubility of 0.6, insolubility was considered a potential drawback for production of rPhaZs. To overcome potential insolubility issues, induction was performed overnight at 15 °C. In addition, extracellular PhaZs are known to be sensitive to dithiothreitol (DTT), suggesting they likely form disulfide bonds [34]; in fact the DiANNA 1.1 web server tool predicted several disulfide bonds for the mature PhaZs used in this study [35-37]. The plasmid pET-22b(+) — previously used to express fusion proteins of the substrate binding domain of PhaZ_*Pst*_ [38], and PhaZs from *P. mendocina* DSWY0601 [19] and *Bacillus* sp. NRRL B-14911 [39] — containing the N-terminal peIB leader sequence for periplasm localization (a more favourable environment for disulfide bond formation) was selected. In addition, we selected the *E. coli* expression strain Rosetta-gami B(DE3) and, as an alternative, T7 Express *lysY/I*^*q*^ (allowing cloning and expression of toxic genes through tight control of expression by *lacl*^*q*^ and of T7 RNA Polymerase by lysozyme). Both strains are BL21 derivates designed to aid in expression of proteins that contain disulfide bonds and suitable for expression under T7 promoter. Furthermore, Rosetta-gami B(DE3) *E. coli* contains the plasmid pRARE that supplies tRNAs for five rare codons (three present in PhaZ_*Cte*_, one in PhaZ_*C*sp_, and two in PhaZ_*Mal*_, PhaZ_*Pst*_, and PhaZ_*R*sp_).

**Table 2.**
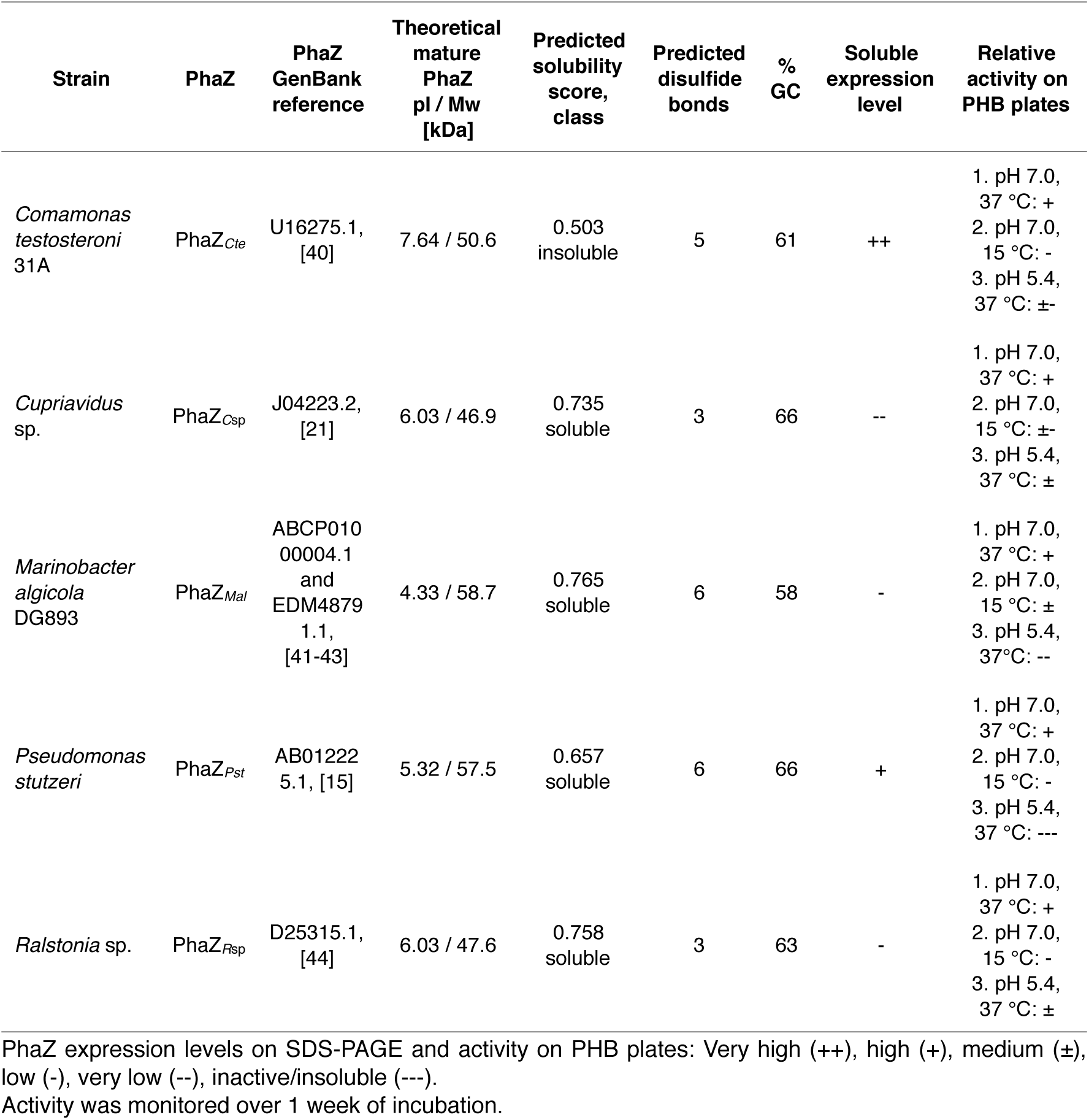
PhaZs properties, sequence analyses, qualitative expression and activity. Relative activity was tested on PHB plates where rPhaZs were deposited and displayed degradation as transparent halos.

Successful pET-22b(+)-PhaZs constructs were obtained for all cloned PhaZs. Sequencing of mature *phaZ* inserts revealed the sequences of PhaZ_*Mal*_ and PhaZ_*Pst*_ were the same as in the Genbank registers, while some changes were found for PhaZ_*Cte*_, PhaZ_*C*sp_, and PhaZ_*R*sp_, that resulted in 3, 1, and 7 amino acid changes, likely due to variations in the taxonomic strains [45] since their deposition.

### 3.2 Expression and purification of rPhaZs

Conditions for induction were established with the expression strain *E. coli* Rosetta-gami B(DE3). SDS-PAGE showed that PhaZ_*Cte*_ and PhaZ_*Pst*_ were present in both SF and IF for inductions with 1 mM IPTG at 15 °C. Decreasing the IPTG concentration reduced insoluble rPhaZs, but the maximum expression in SF was observed at 1 mM. PhaZ_*C*sp_ expression was limited in both SF and IF, while PhaZ_*R*sp_ showed higher accumulation in the IF; the same phenomenon was observed for PhaZ_*Mal*_. When inductions were carried at 37 °C, PhaZ_*Cte*_ and PhaZ_*Mal*_ were only present in the IFs.

Activity was verified for SFs in PHB plates incubated at 30 °C (example shown in Figure 1 (i) for PhaZ_*Mal*_, for which clear zones were only observed for soluble fractions from cultures induced at 15 °C with 0.05 mM IPTG for T7 Express *lysY/I*^*q*^ *E. coli*, and 0.4 or 1 mM IPTG for *E. coli* Rosetta-gami B(DE3)). T7 Express *lysY/I*^*q*^ *E. coli* was tested with PhaZ_*Mal*_ to see if low expression could be due to gene toxicity, but expression was not improved and active enzymes could not be produced with induction above 0.05 mM IPTG even at 15 °C. The ring effect observed in some samples was likely due to self-inhibition at high concentrations of rPhaZ [46-48]. The combined SDS-PAGE/PHB plates analysis is important because rPhaZs at low expression levels were not always observed by SDS-PAGE — the PHB plate assay helps confirm the presence of active rPhaZs in the SFs. For the other rPhaZs, SFs only showed activity when cultures were induced at 15 °C. The favored induction conditions were determined to be 15 °C with 1 mM IPTG for PhaZ_*Cte*_, PhaZ_*C*sp_, PhaZ_*R*sp_, and PhaZ_*Pst*_ and 0.05 mM IPTG for PhaZ_*Mal*_.

**Figure 1.**
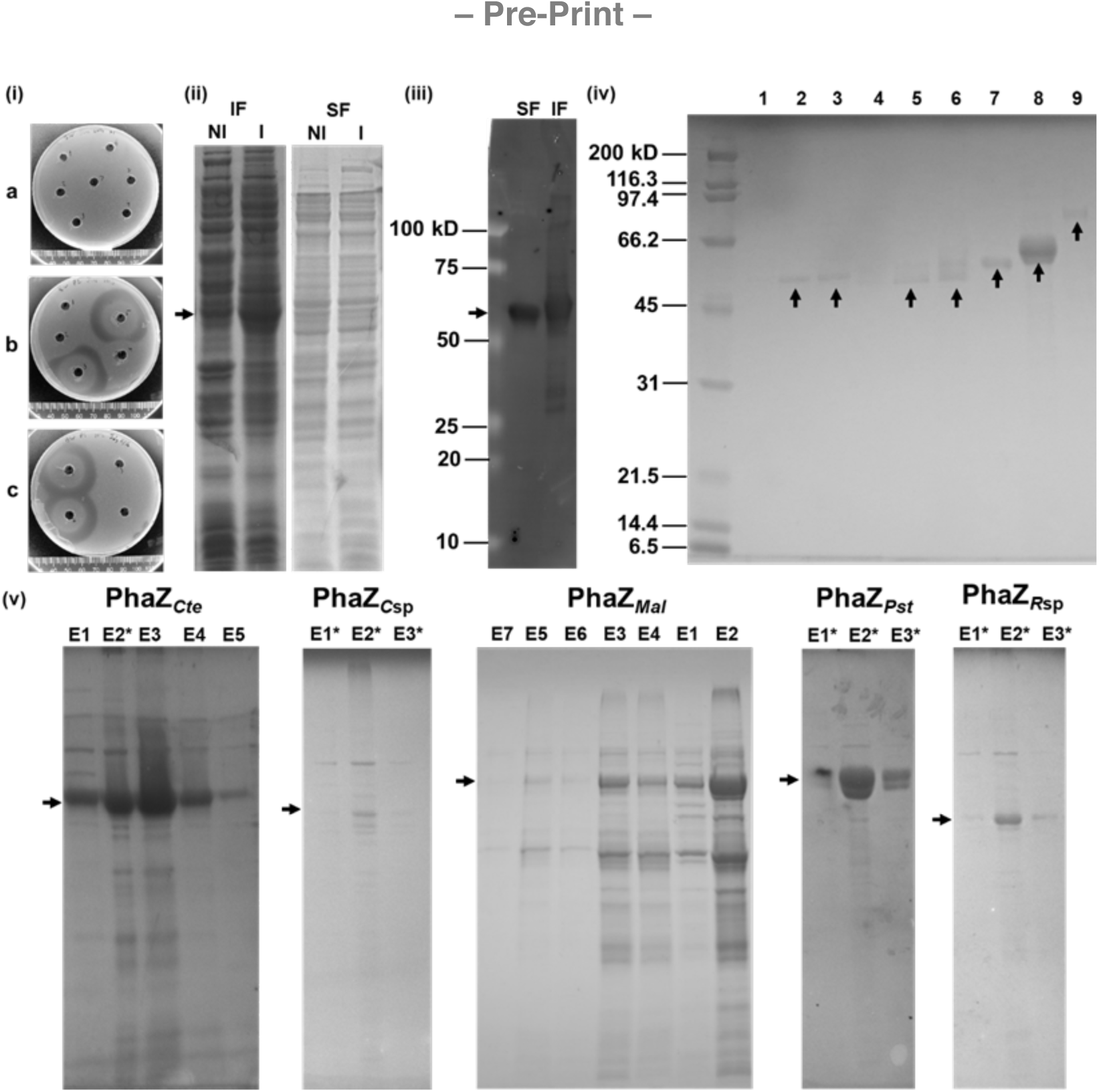
Examples of assays for production of recombinant PhaZs. (i) Induction screening of PhaZ_*Mal*_ on PHB plates (SFs, 30 °C, 7 days incubation): (a) and (b) T7 Express *lysY/I*^*q*^ *E. coli* (induced with IPTG 0.05–1 mM, at 37 °C for 2 h and 15 °C overnight, respectively, in duplicates), and (c) *E. coli* Rosetta-gami B(DE3) (induced with IPTG 0.01–1 mM, at 15 °C overnight). (ii) T7 Express *lysY/I*^*q*^ *E. coli* with construct containing PhaZ_*Cte*_ with signal peptide, induction with 0.1 mM IPTG, at 15 °C overnight (NI: no IPTG added, I: IPTG added; 2 *µ*l loaded for IF and 14 *µ*l for SF). (iii) Western blot of SF and IF of rPhaZ_*Cte*_ containing a His tag (induced with 1 mM IPTG, at 15 °C overnight). (iv) Purified rPhaZs: lanes 1 to 9 are respectively: PhaZ_*C*sp_ F, PhaZ_*C*sp_ C, PhaZ_*C*sp_, PhaZ_*R*sp_ F, PhaZ_*R*sp_ C, PhaZ_*R*sp_, PhaZ_*Cte*_, PhaZ_*Pst*_, PhaZ_*Mal*_ C; where F: filtrate and C: concentrate from size exclusion columns. (v) SDS-PAGE of rPhaZs after imidazole elution (E: consecutive elution; * indicates a 10-fold reduction factor in the loading volume of the elution fraction). Equilibration, sample application, and wash contained 20 mM imidazole, while the elution buffer contained 500 mM imidazole, except for PhaZ_*Mal*_, for which 150 mM were used for E1 and E2, 100 mM for E3 and E4, 200 mM for E5 and E6, and 500 mM for E7. Arrows point to the location of rPhaZs.

Removing the native signal peptide from the PhaZ sequence is a key step in diminishing the formation of inclusion bodies and avoiding completely insoluble PhaZs when using pET-22b(+) or plasmids that add signal sequences. As an example, Figure 1 (ii) shows SDS-PAGE for an induction of T7 Express *lysY/I*^*q*^ *E. coli* with a construct containing PhaZ_*Cte*_ with signal peptide; cultures were induced with 0.1 mM IPTG and incubation was done at 15 °C overnight. Protein production is clearly visible but only observed in the induced IF; PhaZ was not observed in the SF samples, suggesting the formation of inclusion bodies. Induction at higher IPTG concentrations and addition of ethanol in the induction stage [49] did not lead to improved expression.

The presence of the C-terminal His-tag was verified through Western Blot for SF and IF of PhaZ_*Cte*_ and PhaZ_*Mal*_ before proceeding to large-scale inductions for purification. An example is shown in Figure 1 (iii) for the SF and IF of PhaZ_*Cte*_ induced at 15 °C with 1 mM IPTG. After 1-L inductions, all rPhaZs could be separated through simple His-tag based purification (as confirmed by SDS-PAGE in Figure 1 (iv), (v)). Relative expression levels in the SF of each PhaZ can be observed in Figure 1 (v). These were qualitatively classified as very high for PhaZ_*Cte*_, high for PhaZ_*Pst*_, low for PhaZ_*Mal*_ and PhaZ_*R*sp_, and very low for PhaZ_*C*sp_ (Table 2). Purification, which was especially challenging for PhaZ_*Mal*_, could be further improved using a combination of strategies, including doing the equilibration, sample application, and wash steps with solutions containing 50 mM imidazole — this improved purity of PhaZ_*Cte*_, PhaZ_*Mal*_, and PhaZ_*Rsp*_, at the expense of recovery — adding an elution step with 150 mM imidazole instead of 500 mM for PhaZ_*Mal*_, and using size exclusion columns.

### 3.3 Comparison of rPhaZ activity

While PHB plates have been mostly used to screen for PHB degrading bacteria [1], activity from expressed PhaZs has also been estimated by the diameter of clear zones on glass slides covered by PHB-agar mix [9,13] (this test is limited to short-term incubations due to agar drying, but is advantageous for preliminary assessment and when only small volumes of sample are available).

In this study, a rapid method using PHB plates was used to compare PhaZ activity at various pH and temperatures and provided semi-quantitative assessments of activity based on the diameter of degradation halos formed (Figure 2). At 37 °C, degradation was observed on the first day of incubation for all rPhaZs tested, while longer incubation periods were required at 15 °C. PhaZ_*Mal*_ showed the highest activity at 15 °C (halo observed after 1 day) compared to the other rPhaZs (halos observed after 6 days), which could be explained by its marine origin [41]. All enzymes were rendered inactive at pH 4.3 (no halos discernable) but PhaZ_*C*sp_ and PhaZ_*R*sp_ retained significant activity at pH 5.4. This is consistent with their broad pH working ranges (PhaZ_*C*sp_ is stable when stored at pH 5.0–8.0 [50] with optimum activity at pH 7.5 [51], and the optimum pH range of PhaZ_*R*sp_ is 5.0–6.0 [52]).

**Figure 2.**
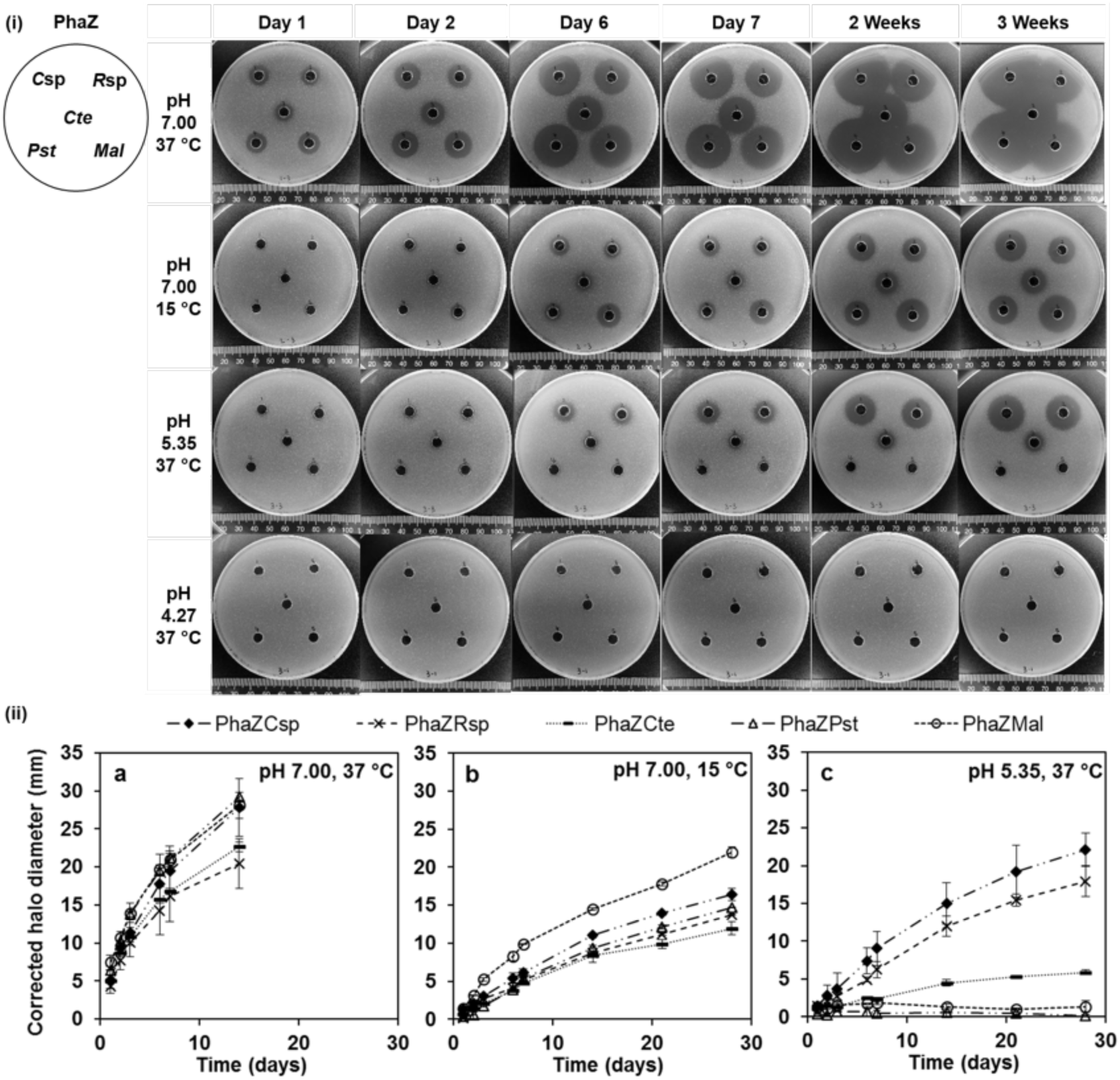
Degradation activity of rPhaZs on PHB plates. rPhaZs concentration was ∼ 2 *µ*g/ml. (i) Degradation as a function of time under different temperatures and pH values. Halos indicate degradation. (ii) Degradation halos diameter (corrected for well diameter). (a) pH 7.00, 37 °C; (b) pH 7.00, 15 °C; and (c) pH 5.35, 37 °C. No halos were observed at pH 4.27, 37 °C.

These results could be confirmed and semi-quantified by comparing the rate of change of the degradation halos under the different conditions tested (Figure 2 (ii))). For example, similar degradation rates were observed for all rPhaZs at 37 °C and pH 7.0, but PhaZ_*Mal*_ had a noticeably greater rate at 15 °C and pH 7.0 (leading to ≈ 33% more degradation after 28 h). Such methods represent powerful tools for screening recombinant and engineered PhaZs, as was demonstrated by Hiraishi et al. who used LB plates containing PHB granules, IPTG, and antibiotics to evaluate clear zone activity of PhaZ mutants [23].

## 4 Concluding remarks

This study presents a streamlined platform for the rapid production of rPhaZs. Five active PHB-degrading extracellular PhaZs (PhaZ_*Cte*_, PhaZ_*Pst*_, PhaZ_*C*sp_ PhaZ_*R*sp_, and PhaZ_*Mal*_), originating from bacteria from diverse environments, were successfully produced in the SF of Rosetta-gami B(DE3) *E. coli*. An important aspect of the method requires the removal of the native signal peptide sequence of PhaZ to avoid production of insoluble proteins and inactive enzymes. Expression levels and purity varied for each enzyme — PhaZ_*Cte*_ and PhaZ_*Pst*_ saw highest expression — but they could all be recovered and retained activity. In addition, degradation activity could easily be assessed by determining the diameter of degradation halos in PHB plates. This assay can be done in parallel for the initial screening of PhaZs and conditions for diverse applications. Both the rPhaZ production platform and the modified PHB plates assay are versatile and reliable, and could be employed with other PhaZs reported in the literature or novel ones to be discovered or synthesized.

## Acknowledgement

This work was supported by the Alberta Agriculture and Forestry Strategic Research and Development program and the Natural Sciences and Engineering Research Council of Canada. BW was supported by the Natural Sciences and Engineering Research Council of Canada Undergraduate Student Research Award program.

